# Queens from a unique hyper-dense *Lasius niger* population tolerate pleometrosis better than queens from a ‘normal’ population

**DOI:** 10.1101/2024.07.16.603683

**Authors:** Stanislav Stukalyuk, Tomer Czaczkes

## Abstract

The claustral, monogynous ant *Lasius niger* often founds colonies pleometrotically (with two or more queens), but later aggression from queens or workers can result in the death of all but one queen. Recently, a hyper-dense population of *L. niger* was discovered, showing minimal worker-worker aggression and interconnected colonies. Here, we ask whether queens are more tolerant of conspecifics in a pleometrotic setting. We collected queens directly after a nuptial flight from both the hyper-dense population and a ‘normal’ population, maintained them in pleometrotic groups, and followed queen survival for 227 days. While queens from the ‘normal’ population showed poor survival (under 20% survival after 130 days), resulting in usually one queen per pleometrotic group, 75% of queens from the hyperdense population survived to 227 days. Mortality in the ‘normal’ population was not centered around the emergence of the first workers. While the colonies from the hyper-dense population are all monogynous, this tolerance of pleometrosis may be linked to their apparent intraspecific tolerance and may be a step towards unicoloniality.

## Introduction

Many species of ants (Hymenoptera, Formicidae) establish their colonies by haplometrosis, i.e., independently and at the expense of the resources of a queen. At the same time, pleometrosis, in which several queens participate simultaneously in the founding of a colony, is a fairly common method of colony foundation among ants (Rissing et al., 2000). In *Lasius niger*, pleometrotic colony foundation has been reported in 18% of cases (Sommer & Hölldobler, 1995) and in up to 40% of other ant species (Trunzer et al., 1998). In primary pleometrosis, the foundation is formed by the pleometrotic group of queens; in secondary pleometrosis, the queens are accepted into the adult colony after the nuptial flight (Zakharov, 1991).

Pleometrosis is known not only in ants, but also in thrips (Morris et al., 2002) and termites (Hartke & Rosengaus, 2013). Pleometrosis in ants could be caused by the preference of queens for pre-existing cavities after the nuptial flight, which they use for shelter and as an initial nest (Tschinkel, 1998). In addition, the limitation of nesting sites may be important for forming pleometrotic groups (d’Ettorre et al., 2005). Pleometrotic groups do not form by chance -in experiments, queens were collected in one chamber when they had several to choose from (Aron & Deneubourg, 2020).

If the species is obligately monogynous, as is the case in *L. niger*, fights between queens eventually occur, resulting in only one surviving (Sommer & Hölldobler, 1995). This is where pleometrosis can offer significant benefits. First, colonies founded by multiple queens grow faster than those founded by a single queen, as demonstrated in L. niger (Sommer & Hölldobler, 1992, 1995) and other ant species (Ostwald et al., 2021). Faster colony growth provides quicker access to food resources, which is particularly advantageous when the density of existing ant colonies is high and resources are limited (Boomsma et al., 1982; Shaffer et al., 2016).

The size of the pleometrotic group is also essential - on average, in *Formica podzolica*, groups of 2-4 queens produced more cocoons and had lower mortality than groups of 8-16 queens (Deslippe, 1994; Deslippe & Savolainen, 1995). On average, one queen can produce three times more workers, and the time from egg to adult worker is also reduced by 4 days (8%) in pleometrotic groups compared with haplometrotic groups (Offenberg et al., 2012). This highlights the potential benefits of pleometrosis, as it not only increases colony growth but also enhances survival rates and efficiency in worker production.

Despite the well-studied phenomenon of pleometrosis, several questions remain unanswered. For example, it is not known whether colony population density can influence pleometrosis. Where population densities are high and further founding space is limited, pleometrosis may assist colonies in overgrowing to the point where they can defend themselves from their neighbors.

Recently, a unique population of *L. niger* was discovered. While *L. niger* is reported as monogynous over the vast majority of its range, this population in the vicinity of Vyshneve city (Kyiv Region, Ukraine) exhibits an unusually high density of nest mounds, encompassing c. 16,000 nest mounds (for details see Stukalyuk et al., 2022, 2023). This population shows limited intraspecific aggression among workers from different nests within the same nest complex, with nests being linked by active trails (Stukalyuk et al., in preparation, data available upon request).

Studying differences in pleometrotic founding between this ‘nest complex’ population and queens from the normal ‘solitary mound’ population may be very informative for understanding the factors that favour pleometrotic or solitary founding. By comparing these two populations, we can investigate how nesting type factors influence the success and survival of queens in different founding contexts. This could help elucidate the evolutionary benefits and trade-offs associated with pleometrosis and solitary founding in *L. niger*.

In this study, we addressed the following questions.

1)Do *Lasius niger* queens from the hyper-dense nest complex population survive longer in pleometrotic conditions than queens from ‘solitary nest mound’ populations? We expected the queens from the ‘nest complex’ population to survive longer under pleometrotic conditions.

2)At what number of workers does fighting between queens start, and does their mortality increase? Does queen mortality increase as the number of workers increases? In addition, we asked whether queens are killed or die singly or all in one event.

## Materials and Methods

### Experiments

The *Lasius niger* nest complex B covers an area of 13.3 hectares and contains 15,599 nests with a average density of 11.7 nests per 100 square metres. However, this density may be underestimated as only clearly visible mounds were surveyed. The complex is elongated and irregular in shape. The average diameter of the mounds is 34.1 cm and the average height is 22.9 cm. The density of nests varies according to the zone of the complex B: in zone I the density is 15.9 nests per 100 square metres, in zone II - 11.9, in zone III - 11.5 and in zone IV - 4.4 nests per 100 square metres.

Intensive communication between mounds was observed, including both above-ground trails and underground tunnels connecting main and secondary nests. Auxiliary nests often occur around aphid colonies and can develop into primary nests over time. The average population of a nest with a base mound diameter of 35 cm is about 14,000 workers, and the maximum population in a monogynous nest can reach about 60,000 workers.

The *Lasius niger* nest complexes discovered are unique in that they contain tens of thousands of nests. The study (Stukalyuk et al., 2023) included about 30 such complexes, of which only two were of such impressive size. They differed from individual nests, which were about 12 km apart, and did not form clusters. This rules out the possibility that queens found in nesting complexes could be isolated queens from individual nests. The geographical and spatial separation is also confirmed: the distance between individual nests and nesting complexes is 50 metres. Therefore, the results of our study suggest that factors related to nest type play an important role in the survival of Lasius niger queens in pleometrotic groups.

Laboratory experiments to determine the level of aggression between young queens during pleometrotic founding were performed as follows: Thirty-seven (most probably inseminated) dealate *Lasius niger* gynes were collected from the ground within the territory of nest complex B on 10 July 2019, immediately after the nuptial flight. On the same day, a group of 21 dealate gynes was collected from the ground in a territory with scattered monodomous *L. niger* nests in the Feofaniya Park (50.3408°N, 30.4869°E) in the suburb of Kyiv, about 12 km ESE from the nest complex (solitary nest mound group). The likelihood of mixed groups of queens from different territories within the nest complex is very low, as the majority of dealate queens collected from nest complex B are likely to originate specifically from this complex.

The collected gynes were placed in test tubes 25 cm long and 4 cm in diameter. One third of the tubes were filled with a moist cotton plug. The queens from the nest complex examined (nest complex group) were placed in seven test tubes. Six of the tubes contained five gynes each, and the seventh tube contained seven gynes. The control groups of queens (solitary nest mounds population) were placed in four tubes. Three of the tubes contained five queens each, and one tube contained six gynes. Overall, tubes from the area of the nest complex underwent 298 inspections, and the control tubes 74 inspections.

A difference in the number of observations between groups is due to the fact that groups of queens from solitary nest mounds reduced the number of observations to a one queen compared to another group, and the observations were stopped.

The tubes were stored in the dark at a temperature ranging from 20 to 21°C and inspected every five days at noon with the help of a magnifying glass (magnification 10×). The following variables were recorded: (1) the number of queens alive, (2) the number of queens that probably died of natural causes (those whose dead bodies appeared intact), (3) the number of queens killed (those whose dead bodies showed sign of damage), (4) the number of larvae, (5) the number of pupae.

In most cases, however, the dead queens were at the other end of the test tube and their bodies were intact. In the case of fights, we observed that the bodies of the queens were always dismembered. The queens were also scored for whether clustered or dispersed in the tube. After the first workers emerged, the incipient colonies were fed with sugar syrup, and small fruit flies (*Drosophila* sp.) killed by freezing after each inspection. When all but one queen in the tube died, the tube was excluded from further observation. The experiment was planned to run until the second generation of workers in tubes with queens from nest complex. However, due to the COVID-19 related lockdown, the experiment was forcibly ended on 27 February 2020, approximately 227 days after it began. This was not a crucial constraint, as the deciding criterion, behavior after the eclosion of the first workers, could be fully observed. None of the tubes with *Lasius niger* hibernated. This species is capable of developing without hibernation, as has been shown by numerous observations of fans of ant colonies (myrmekeepers).

The determination of the species of *Lasius niger* was carried out according to the keys given in the work of B. Seifert (2020).

### Statistical analysis

#### Survival analyses

Survival analyses on the two *Lasius* niger populations were conducted using the coxme package (Therneau, 2024) run with the software package R (R Development Core Team 2012) by fitting Kaplan-Meier survival curves, including tube identity as a random effect and performing log-rank tests. The complete code and output used for this analysis are provided in supplement S1. We used ANOVA to compare the number of brood and workers on tubes from queens who died of natural causes and those who died due to fights.

## Results

### Effect of nest type on survival of queens in *Lasius niger*

Queens from the ‘nest complex’ population had a higher survival in pleometrotic groups than queens from the ‘solitary nest mound’ population (Chi-square = 14.6, p=1e-04, Figure 1).

**Figure 1.**
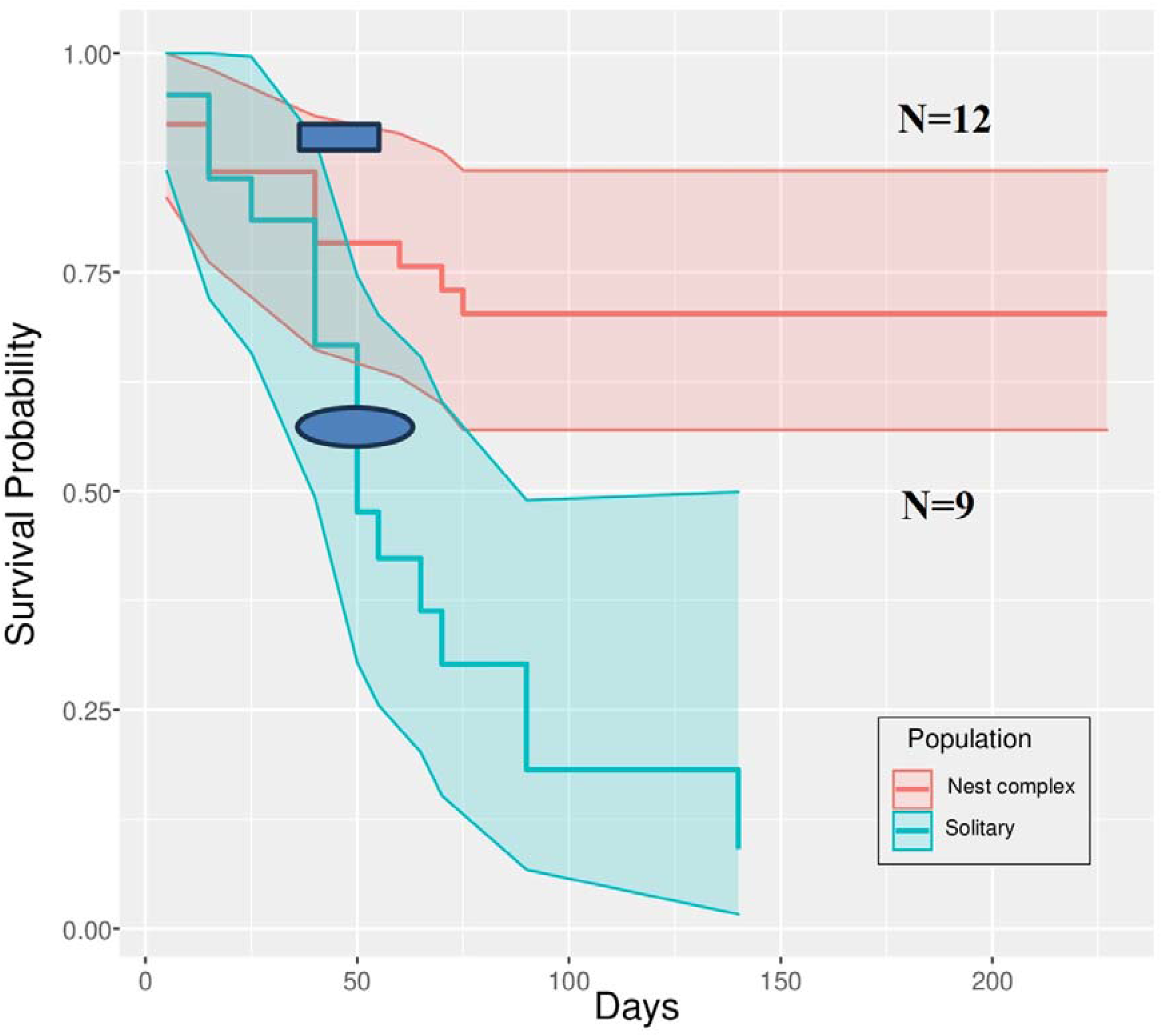
Survival curve for *Lasius niger* queens from a hyper-dense nest complex and a solitary nest mound population. Ribbons are 95% confidence intervals. Rectangle - time of eclosion of the first workers in the group of the nest complex (test tubes 3, 6, 7 - 40 days, test tube 4 - 35 days, test tube 1 - 55 days), oval - in the group of solitary nest mounds (test tube 3 - 35 days, test tube 2 - 40 days, test tube 5 - 65 days).

In the solitary nest mound group, the number of queens decreased to one by the 130th day of observation, whereas in the nest complex group, at least four queens remained alive per group until the end of the experiment.

At the time of the release of the first workers, there were 6, 4 and 4 living queens in three of the four test tubes from the single nest mound group. Within 23 days of the appearance of the first workers, one surviving queen remained in each of these three tubes. In the fourth test tube, the number of queens remained unchanged (5), since the appearance of the brood was not recorded during the observations in this tube, so we did not take it into account.

One third (7/ 21) of queen deaths occurred before the first workers emerged (Figure 2).

**Figure 2.**
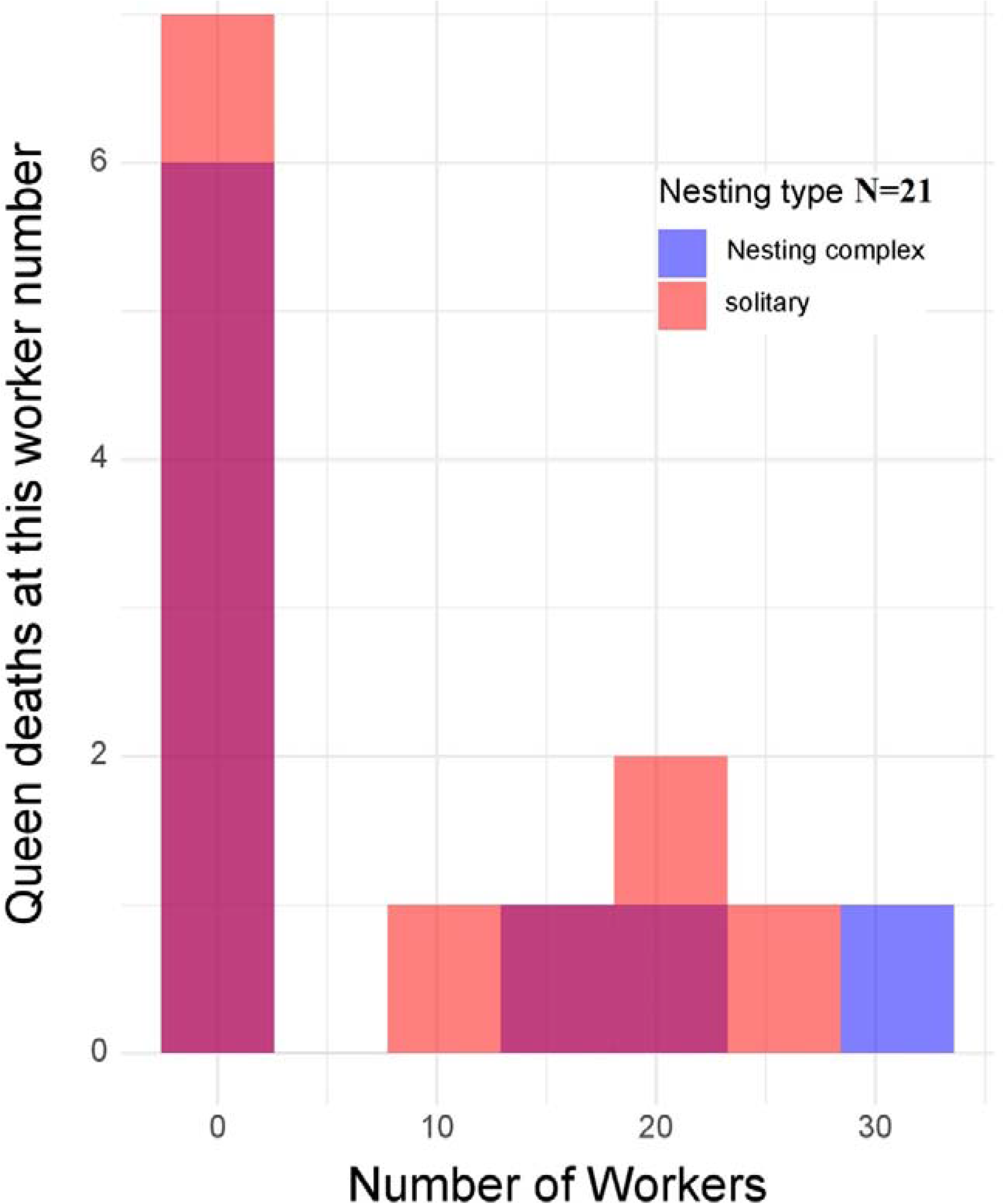
Mortality rates of queens as a function of number of workers and time of study.

The remaining mortalities occurred when a group of 10-30 workers had already emerged from the cocoons. Mortality patterns appear similar between the nest complex and solitary nest mound groups. In most cases, the number of queens decreases by one between observations, much less often by two (average 1.3, Figure 3).

**Figure 3.**
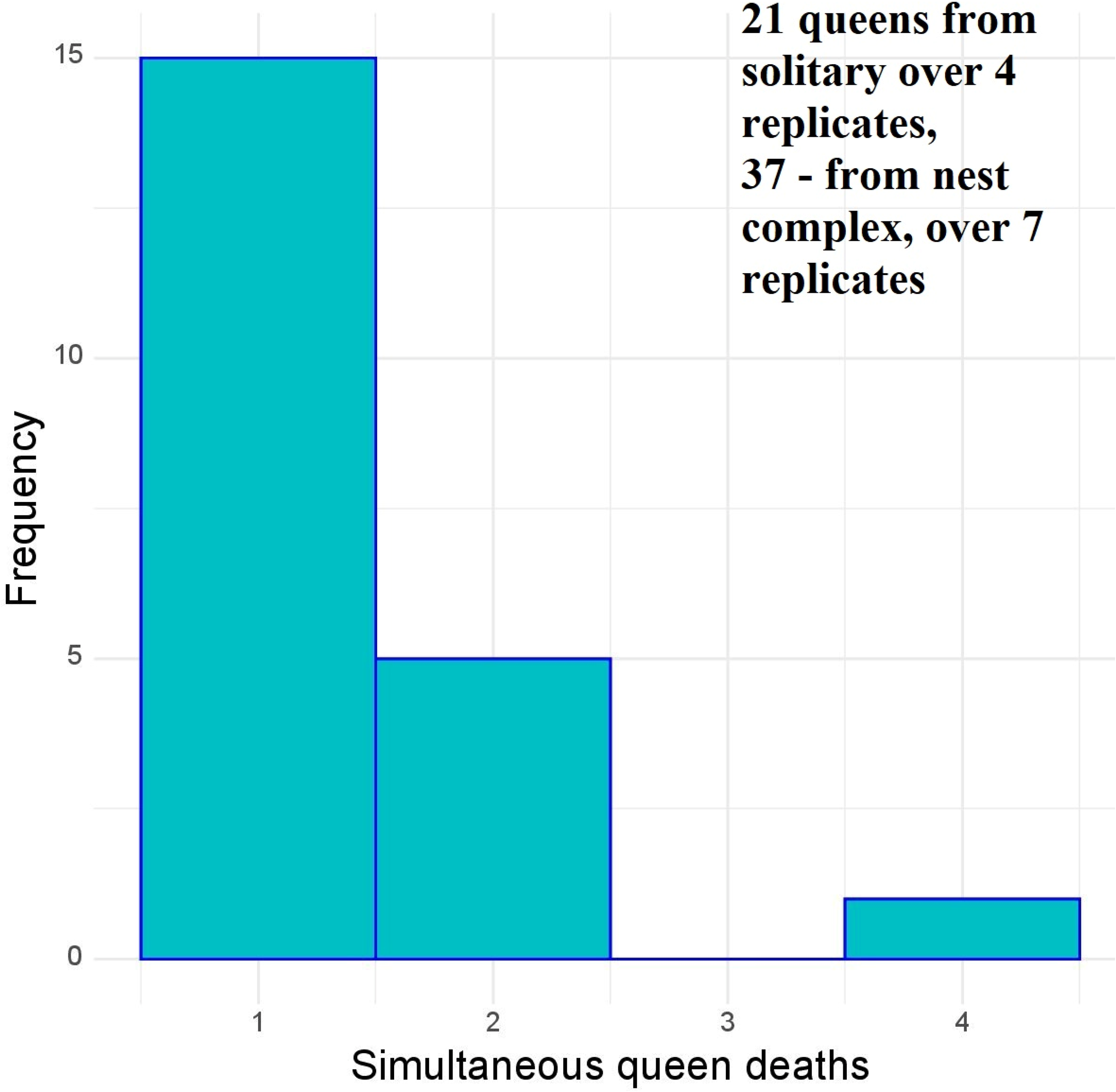
The number of queens dying in one observation period (5 days)

We did not find a significant difference in the number of workers (ANOVA, F=2.918, df=24, p = 0.1079) and pupae (F=0.11, df=24, p = 0.740) found in tubes with queens that died due to fights or those that died of natural causes. However, we note that there were twice as many larvae in tubes when queens died due to fights, than in tubes when queens died of natural causes (F=6.23, df=24, p = 0.021).

In all cases the queens were in the same group, next to the source of moisture (cotton at the end of the test tube separating the chamber with the queens from the water). Therefore, only the number of workers and brood for the whole pleometrotic group could be taken into account at the time of detection of a dead queen.

**Supplementary Table 1.** Indicators of the number of workers and brood at different stages of the life cycle of experimental colonies.

| nesting_type | number of tube | Larvae | puppaes | number_workers | number_queen |
| --- | --- | --- | --- | --- | --- |
| nest_complex | uk6, before workers eclosion | 14 | 4 | 0 | 4 |
|  | Last observation | 23 | 0 | 32 | 4 |
|  | uk7, before workers eclosion | 43 | 15 | 0 | 7 |
|  | Last observation | 55 | 0 | 52 | 7 |
|  | uk4, before workers eclosion | 20 | 35 | 0 | 5 |
|  | Last observation | 54 | 0 | 51 | 4 |
|  | uk3, before workers eclosion | 10 | 21 | 0 | 4 |
|  | after reduction of queens to 1 | 26 | 5 | 31 | 1 |
|  | uk2, before workers eclosion | 15 | 0 | 0 | 4 |
|  | Last observation | 0 | 0 | 0 | 3 |
|  | uk1, before workers eclosion | 17 | 10 | 0 | 4 |
|  | Last observation | 35 | 0 | 39 | 4 |
|  | uk5, before workers eclosion | 8 | 0 | 0 | 4 |
|  | after reduction of queens to 2 | 0 | 0 | 0 | 2 |
| Average number per type of nesting | before workers eclosion | 18.1±4.4 | 12.1±4.8 | 0 | 4.5±0.4 |
|  | Last observation | 27.5±8.5 | 0.7±0.7 | 29.2±8.1 | 3.5±0.7 |
| Solitary nest mounds | f2, before workers eclosion | 25 | 20 | 0 | 5 |
|  | after reduction of queens to 1 | 37 | 13 | 16 | 1 |
|  | f3, before workers eclosion | 35 | 24 | 0 | 4 |
|  | after reduction of queens to 1 | 40 | 15 | 24 | 1 |
|  | f4, before workers eclosion | 12 | 0 | 0 | 3 |
|  | after reduction of queens to 1 | 0 | 0 | 0 | 1 |
|  | f5, before workers eclosion | 23 | 4 | 0 | 4 |
|  | after reduction of queens to 1 | 23 | 0 | 20 | 1 |
| Average number per type of nesting | before workers eclosion | 23.7±4.7 | 12.0±5.8 | 0 | 4.0±0.4 |
|  | after reduction of queens to 1 | 25.0±9.1 | 7.0±4.0 | 15.0±5.2 | 1.0±0 |

From the data in the Supplementary Table 1 it is clear that the type of nesting has a significant effect on the dynamics of the number of workers, larvae and pupae, with the complex nest group showing higher survival and productivity of workers compared to the solitary nest mound group. The number of larvae at the time of the second observation is comparable in the two groups, the number of pupae is 10 times higher in the solitary nest mound group, the number of workers is 2 times higher in the nest complex group. There are differences in the number of queens, workers and brood between different tubes within the same group (Figure 4). For example, in test tube uk3 the number of queens decreased to 1, possibly because queens from the nest complex and from solitary nest mounds could enter this test tube. In the remaining tubes from the nest complex group the number of queens did not decrease, except in uk5 where no brood or workers were observed (Figure 4). In the group of solitary nest mounds, the number of queens decreased to 1 in all cases, but in test tube f4 there was only one queen left at the time of the last observation, with no brood and no workers (Figure 4). This indicates that in some cases a pleometric group of queens may not produce workers or brood due to the influence of various factors.

**Figure 4.**
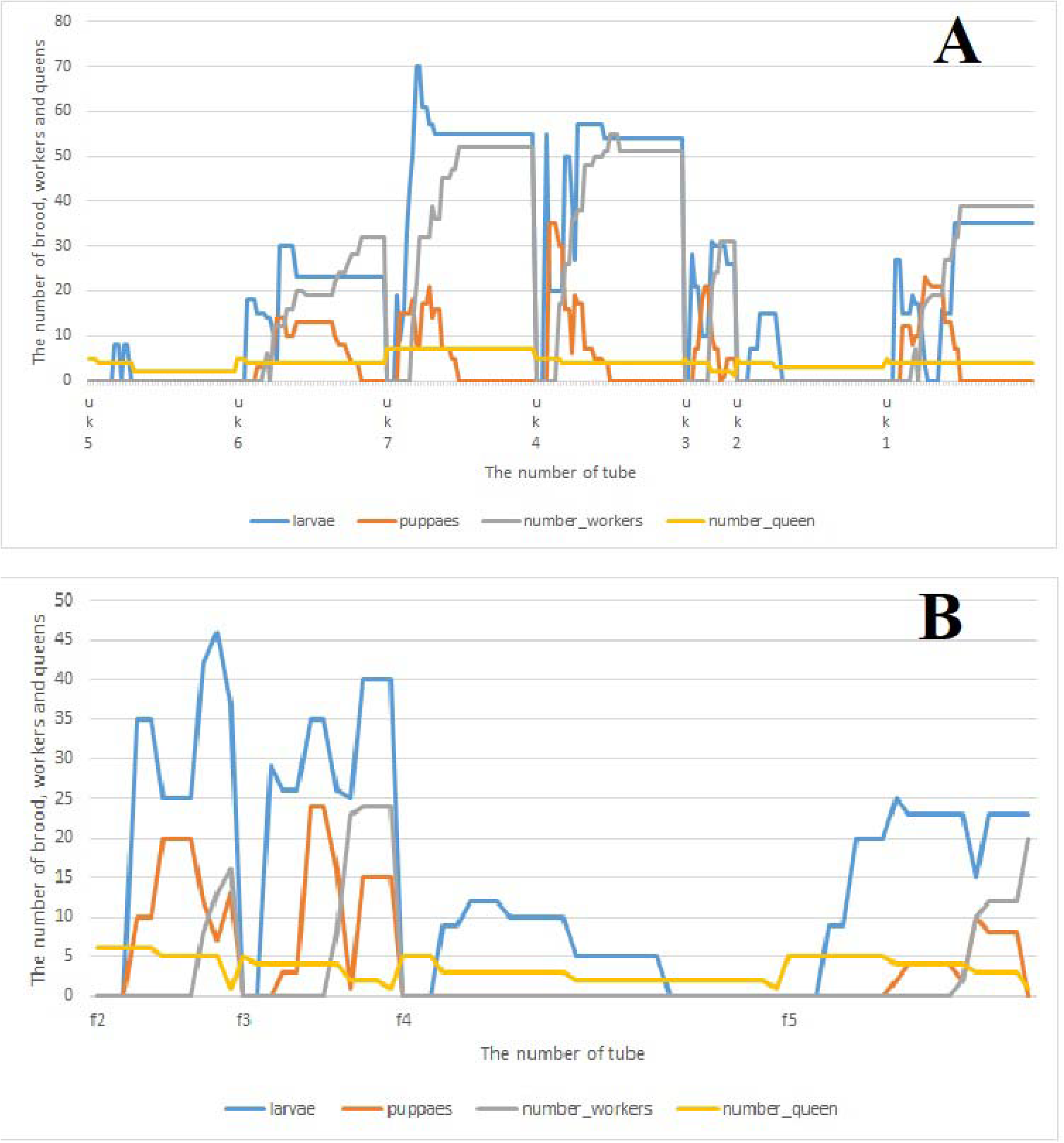
Dynamics of changes in the number of brood, workers and queens for test tubes from the group of the nest complex (A) and solitary nest mounds (B).

## Discussion

The aggressiveness of ants towards strangers (own and other species) is determined by differences in the ant cuticle hydrocarbon profile. For example, the cuticular hydrocarbons (CHC) profiles of *Aphaenogaster iberica* ants vary with altitude and temperature, affecting the recognition of their own and strangers in the colony (Villalta et al., 2020). These differences in CHC profiles are associated with levels of aggression between ants from different populations, indicating the importance of climatic conditions in the formation of the colony signature.

In *Lasius* ants (*L. niger, L. platythorax*), adaptation to different climatic conditions leads to an increase in aggression and chemical differences between former relatives (Wittke et al., 2022).

Only a small subset of CHCs differs between colonies during adaptation, and a few compounds explain individual aggression, suggesting an over-encoding of colony identity. The contribution of individual CHCs to colony differences and adaptation changes are negatively correlated, indicating partial functional separation. The functions of CHCs in insects are so interrelated that ants cannot optimise all of them at the same time. The main problem is the need to maintain a certain physical state of the CHC layer, which depends on its composition and affects its functioning. Also, the need to separate CHC functions may depend on the environmental conditions and life history of a particular species. It is also known that the eggs and larvae of the early stages have individual CHC signatures (de Fouchier et al., 2023). It was found that Camponotus floridanus ants can adapt their recognition of kin by taking into account changes in the chemical profile of the colony (Neupert et al., 2018). These results suggest that ants have multiple templates for nestmate recognition.

Another study hypothesised that no individual ant knows the chemical identity of the entire colony; instead, the identity of the colony is determined collectively by all the ants (Esponda & Gordon, 2015). Each ant responds to the chemical profile of other ants based on its own experience with them. Thus, each ant recognises a particular set of chemical profiles as those of intruders. This model predicts that the results of behavioural tests can be variable, depending on the number of ants tested, and can also change over time and with the experience of individual ants. Such a distributed system allows the colony to identify intruders without requiring all individuals to have the same complete information, which also makes it easier to track changes in chemical profiles because only a subset of ants need to respond to ensure an adequate response.

Czaczkes et al., (2024) found that overt aggression did not vary with level of relatedness or spatial distance in Lasius niger, but differences in antennation and jerking were found between individuals with different levels of relatedness and spatial distance. These observations suggest that ants exhibit different types of responses that may mitigate the need for overt aggression.

Although our study does not directly relate survival to cuticular hydrocarbon composition, our results indicate a significant influence of intrinsic factors, such as nest type, on queen survival in pleometrotic groups.

Our data suggests a higher numbers of larvae during queen death due to fights per tube, contrasting with queens dying of natural causes. Our findings support the notion that queen mortality primarily occurs during the early stages of colony founding.

Our finding of the long-term coexistence of several queens within a pleometrotic group is not the first. As early as 1979, V. E. Kipyatkov noted the long-term (more than 1.5 years) coexistence of such *Lasius niger* queens with a large number of workers (Kypiatkov, 1979). The author notes that, in this case, the transition to obligate monogyny could be very long. Our data confirms this for 227 days.

Especially interesting to us is the very large difference between the ‘normal’ (solitary nest mound) and hyper-dense’ nest complex’ populations, given that this species is obligately monogynous. Under natural conditions, pleometrotic groups typically contain between 2 and 4 queens, with a maximum of 9 (Bartz & Hölldobler, 1982), so the group size we used (5) is realistic and close to typical.

Hyper-dense *L. niger* nest complexes are common in Ukraine (Stukalyuk et al., 2022; 2023), as well as in the European part of Russia (Zakharov, 2015). Therefore, it is likely that the nest complex we studied is not unique; it was simply the first to be studied in detail (Stukalyuk et al., 2022, 2023). Unclear is whether other hyper-dense nest complexes also show strong interconnectedness and low intraspecific aggression to members of the nest complex. Under densely populated conditions, pleometrosis is an effective method of establishing new colonies. Given such a dense population, the greater tolerance of pleometrosis we demonstrated should allow newly founded colonies to establish a competitive colony with shared resources quickly. Given the obligate monogyny of *L. niger*, sooner or later, only one queen remains in all such pleometrotic groups. The duration of the transition to monogyny remains open. Still, our results suggest that this period is much longer in pleometrotic groups from nesting complexes than in solitary nest mounds, and could potentially last for several years.

Fights between L. niger queens can lead to death in pleometrotic groups in 60% of cases (Madsen & Offenberg, 2017), which was confirmed by our experiment, where the number of queens was reduced to one in 3 out of 4 tubes from the group of single nest mounds.

Interestingly, pleometrotic queens have higher survival rates than haplometrotic queens (Bernasconi & Keller, 1996, 1999). The next peak in queen mortality occurs when workers have emerged from their cocoons (Sommer & Hölldobler, 1995). In our case, the number of larvae in groups with queens that died in fights was also two times higher than in groups with queens that died of natural causes, again supporting this dual peak in mortality: a first peak due to natural causes directly after founding, and a second peak after the emergence of the workers.

*Lasius niger* queens investing in high reproduction may experience decreased survival and vice versa (Pamminger et al., 2016). This may be the reason for the high survival rate of queens, at least in the group of the nest complex.

Tolerance between queens is key to the possibility of pleometrosis (Overson et al., 2014). We show that the nesting typical for a given ant species in a given part of its range strongly influences tolerance between queens. Another interesting finding of our study is the long transition period from the pleometrotic group to monogyny. However, at the end of the experiment, there were at least four queens in the nest complex groups.

The *L. niger* queen ants collected from the nested complex mostly survived when kept in pleometrotic groups, while most queens from the normal single nest mound population did not. What could the mechanism behind this difference be? As the pleometrotic queens from nest complex would not be tolerated by non-pleometrotic’normal’ queens, distinct geographical ranges may appear despite mating flights allowing the queens to disperse. This hypothesis would predict that other separate populations of pleometrotic *L. niger* may emerge or already exist.

Alternatively, pleometrotic tolerance may be ontological in origin, either due to developmental changes when developing in a pleometrotic nest or by the virgin queen learning to tolerate a range of workers with reduced relatedness in their natal nest. Uncovering the mechanism behind this pleometrotic tolerance would provide valuable insights into the evolution of pleometrosis. These hyper-dense nest complex populations also offer a range of other opportunities. Given that workers from this population are also less aggressive to non-nestmates from this population (Stukalyuk et al., in prep), the mechanisms enforcing heterometrosis may be the same as those involved in nestmate recognition.

Our study has several limitations. 1. Each group of queens was confined to a test tube and had no opportunity to disperse. The only option was to disperse along the length of the test tube, but all the queens were always grouped next to the cotton, behind which was a supply of water. In several cases, queens at the exit of the test tube were later found to have died of natural causes. The test tubes were separated from each other, so that queens from different groups were separated from each other. As a result of the coexistence of queens and their brood, the profile of cuticular hydrocarbons may have been homogenised, resulting in a decrease in aggressiveness between queens and between workers. We did not observe any fighting between workers.

1. Another important point is the small sample size. Although the effect sizes observed are very large, repeating the experiment with a larger sample size would be ideal.
2. Some of the queens that died of natural causes may have died as a result of stress – it is impossible to distinguish the two.
3. Another limitation of this study may be that the ant colonies were not overwintered. Overwintering is a prerequisite for colony reactivation in *L. niger* (Kipyatkov, 1995); it was not present under the experimental conditions, but colony growth continued. It may be necessary to repeat the experiment with a larger sample and with the obligatory presence of hibernation.

## Conclusions

*Lasius niger* queens, a hyper-dense, low-aggression nest complex group, survived well under pleometrotic conditions, with at least four queens in each group remaining alive until the end of the experiment. By contrast, *L. niger* queens from the solitary nest group had low survival, with the number of queens reduced to 1 in each replicate by day 130. Mortality from both populations had two peaks: a major peak before worker emergence and another one after at least ten workers had emerged. The first peak may represent weaker queens dying from natural causes, while the second may be due to aggression from workers or the other queens.

Taken together, the results of this study demonstrate that factors intrinsic to the queens, not external environmental factors post-pleometrotic founding, drive survival in pleometrotic groups. However, whether these innate or developmental factors are genetic is unknown.

## Supporting information

https://drive.google.com/file/d/1DNjoiqqbP0bBeYGwXEadfBy4EeT7Vxuo/view?usp=drive_link

## Acknowledgements

The authors are grateful to the Editor and to two anonymous reviewers whose comments and suggestions made it possible to improve this paper considerably. The authors are grateful to B. Seifert for help in determining the species of the collected ants *Lasius niger*.

