## Supplementary material for "Queens from a unique hyper-dense *Lasius niger* population tolerate pleometrosis better than queens from a ‘normal’ population": https://drive.google.com/file/d/1DNjoiqqbP0bBeYGwXEadfBy4EeT7Vxuo/view?usp=drive_link: pleometrosis_analysis_for_supplement.pdf

### Pleometrosis in Lasius niger

Tomer J Czaczkes

2024-06-30

#### Contents

**Packages** 1

**Import data** 2

**survival analysis - classic** 2

survival analysis with random effects: . . . . 4

survival analysis - ggplot . . . . 4

**Lasius niger queen death - happens with workers or without?** 5

**Is simultaneous queen killing common?** 6

**Session information - R version, packages, etc** 7

This is the code and output supplement to "Queens from a unique hyper-dense Lasius niger population tolerate pleometrosis better than queens from a 'normal' population" by Stukalyuk and Czaczkes

In this study, queens from a hyper-dense, interconnected population of Lasius niger, and queens from a 'normal' population, were housed together in groups of 5. Survival was recorded regularly, along with other variables such as time of queen death, number of workers, etc.

#### Packages

here we will import all the packages we need. Also will set the seed, to make the analysis repeatable

```
library(readxl) # this is for importing excel files
library (ggplot2) # for making figures
library(glmmTMB) # for stats
library (car) # for Anova function
```

```
## Loading required package: carData
```

```
library (DHARMa)
```

```
## This is DHARMa 0.4.6. For overview type '?DHARMa'. For recent changes, type news(package = 'DHARMa')
```

```
library(emmeans)
library (survival)
library(coxme)
```

```
## Loading required package: bdsmatrix

##
## Attaching package: 'bdsmatrix'

## The following object is masked from 'package:base':
##
##     backsolve
```

```
library(ggfortify) # for turning survival analysis into a dataframe for ggplot
library (eoffice) # fpr exporting to ppt

set.seed(123)
```

### Import data

Note that data on other species was also collected, but will not be presented in this study

### survival analysis - classic

Here we carry out a standard survival analysis using Kaplan-meier curves

```
# Fit Kaplan-Meier survival curves


survfit_obj <- survfit(Surv(day_of_death, died_by_end_of_experiment) ~ nesting_type, data = data)

# Plot the Kaplan-Meier curves for groups defined by nesting_type
plot(survfit_obj, col = c("blue", "red"), lwd = 4, xlab = "Time", ylab = "Survival Probability", main =
```

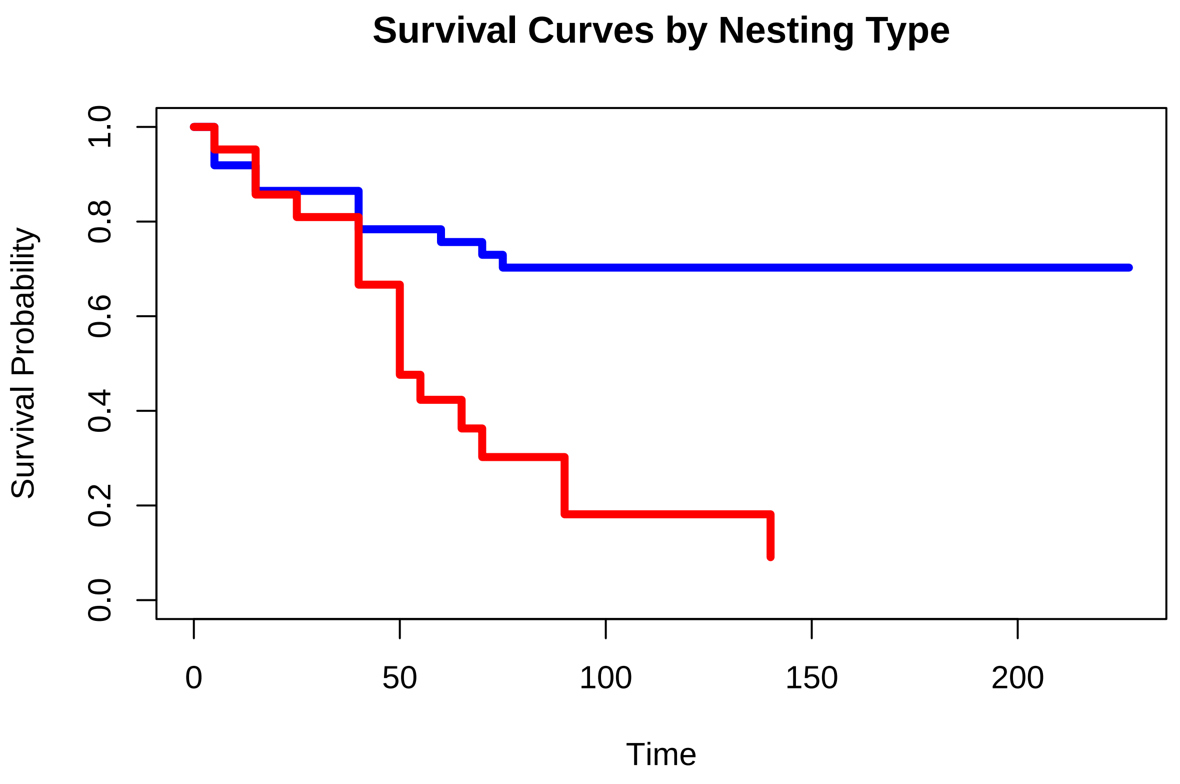


```r
# Add individual experiment survival curves
exp_ids <- unique(data$experiment_ID)  # Get unique experiment IDs
```

```
## Warning: Unknown or uninitialised column: `experiment_ID`.
```

```r
for (id in exp_ids) {
  exp_data <- subset(data, exp_ID == id)  # Subset data for each experiment
  survfit_exp <- survfit(Surv(time = days_since_start, event = number_queen > 0) ~ nesting_type, data =
  lines(survfit_exp,  col = c("blue", "red"), lty = 2)  # Add lines for each experiment
}

# Perform log-rank test
survdiff_obj <- survdiff(Surv(time = day_of_death, event = died_by_end_of_experiment) ~ nesting_type, da
survdiff_obj
```

```
## Call:
## survdiff(formula = Surv(time = day_of_death, event = died_by_end_of_experiment) ~
##     nesting_type, data = data)
##
##                              N Observed Expected (O-E)^2/E (O-E)^2/V
## nesting_type=nest_complex 37       11    19.78      3.90      14.6
## nesting_type=solitary     21       17     8.22      9.37      14.6
##
##  Chisq= 14.6  on 1 degrees of freedom, p= 1e-04
```

Big difference between the nesting complexes. Queens from the nest complex population survive much better in groups than those from the solitary normal population.

##### survival analysis with random effects:

```
# Fit a Cox proportional hazards model with a random effect for experiment_ID
coxme_obj <- coxme(Surv(day_of_death, died_by_end_of_experiment) ~ nesting_type + (1 | experimentID), da

# Print the summary of the model
summary(coxme_obj)
```

```
## Mixed effects coxme model
##  Formula: Surv(day_of_death, died_by_end_of_experiment) ~ nesting_type +      (1 | experimentID)
##     Data: data
##
##   events, n = 28, 58
##
## Random effects:
##          group  variable        sd   variance
## 1 experimentID Intercept 0.1001339 0.01002681
##                   Chisq   df         p   AIC  BIC
## Integrated loglik 13.01 2.00 0.0014928  9.01 6.35
##  Penalized loglik 13.49 1.21 0.0003542 11.08 9.47
##
## Fixed effects:
##                       coef exp(coef) se(coef)    z        p
## nesting_typesolitary 1.421     4.139    0.405 3.51 0.000453
```

##### survival analysis - ggplot

Using GGplot allows us to modify the figure and get confidence interval ribbons

```
plot (plot1)
```

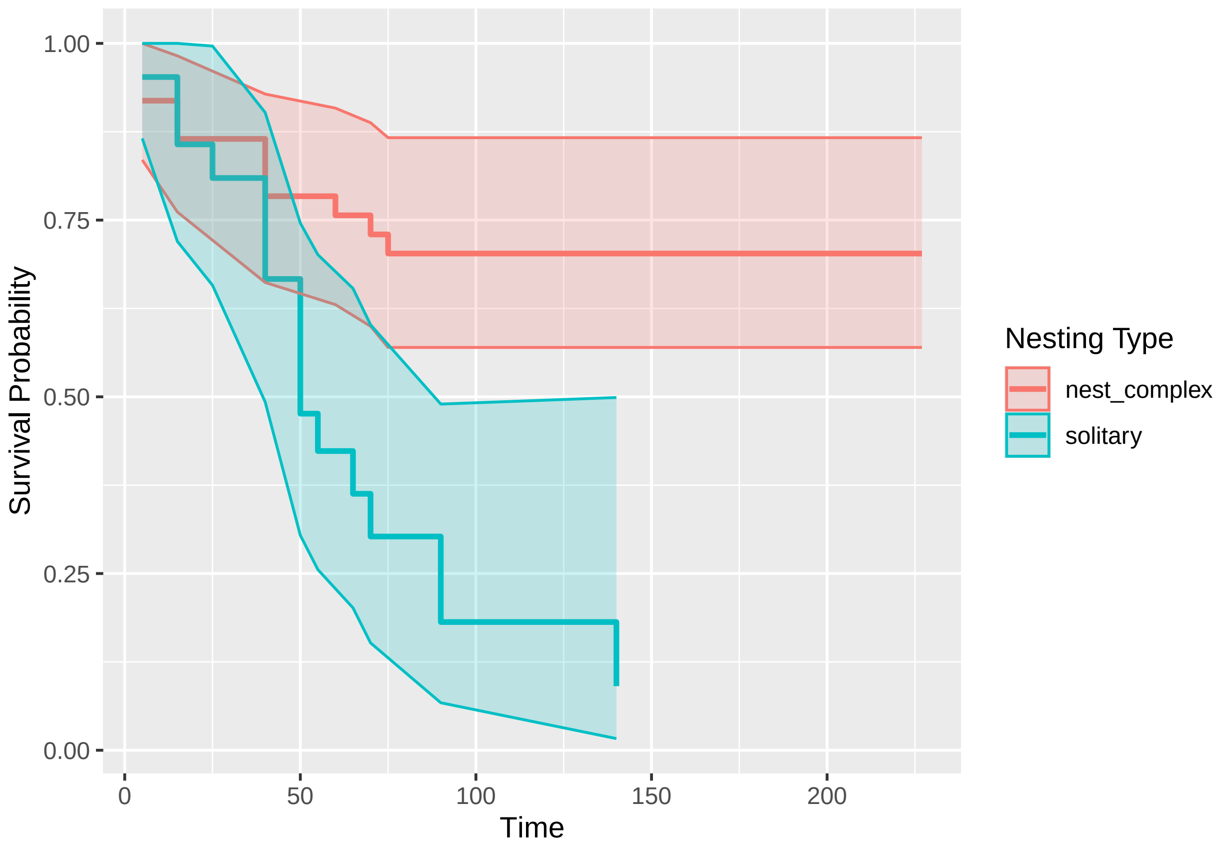


#### Lasius niger queen death - happens with workers or without?

Does queen death happen mostly when workers are present, or before?

First import relevant data

```
workerdata <- read_excel("Data_Experiment.xlsx",
    sheet = "lnigerclean")

#only dates when a queen died
onlyqueendeath <- subset(workerdata, queen_death_event == 1)
```

now a histogram of counts when queens died

```
plot2 <- ggplot(onlyqueendeath, aes(x = number_workers, fill = nesting_type)) +
  geom_histogram(position = "identity", alpha = 0.5, bins = 7) +
  labs(x = "Number of Workers", y = "Frequency", title = "Histogram of Number of Workers by Nesting Typ
  scale_fill_manual(values = c("blue", "red")) +
  theme_minimal()

plot2
```

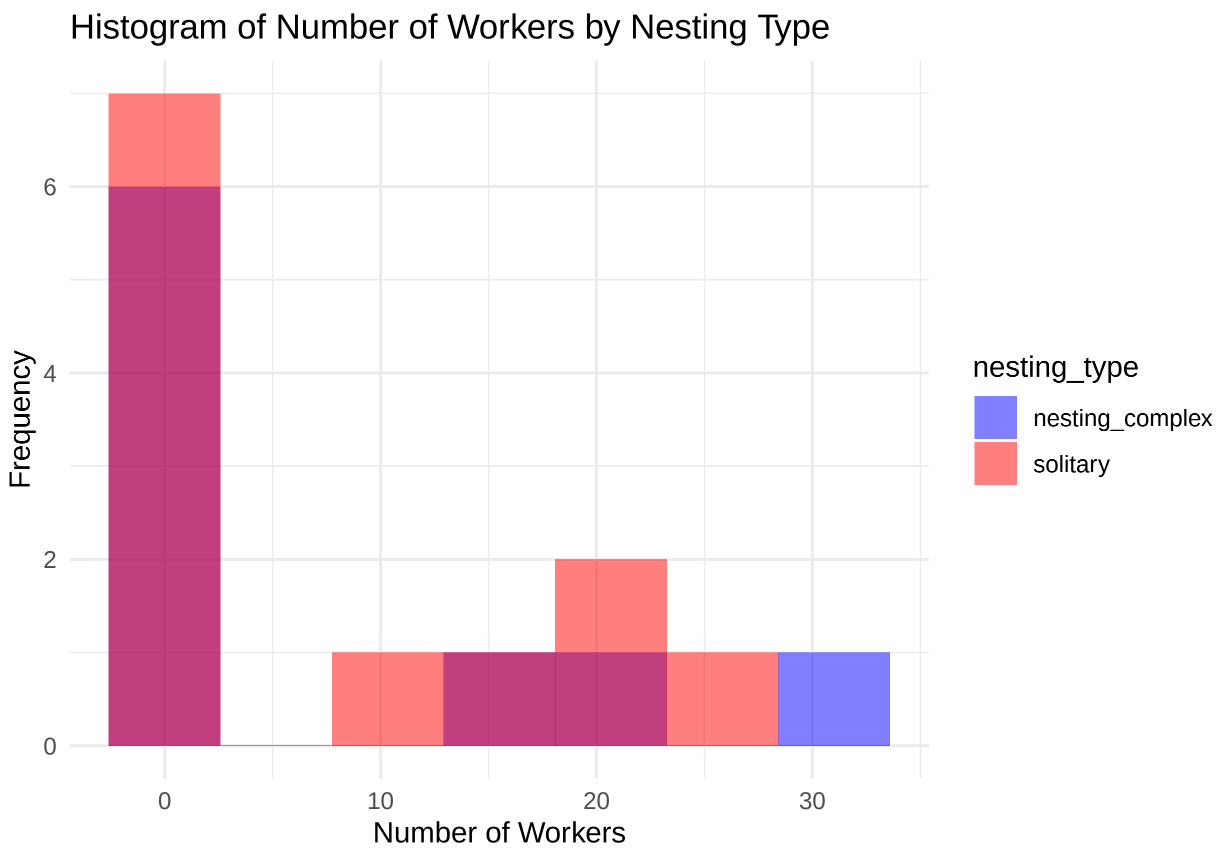


Please note we conduct no formal statistical analysis - the data is too sparse

#### Is simultaneous queen killing common?

Here we ask if queens often die simultaneously in one observation window, or singly. Again, this is only descriptive, not a formal analysis.

```
plot3 <- ggplot(onlyqueendeath, aes(x = killed_in_one_go)) +
  geom_histogram(position = "identity", bins = 4, fill = "#00BFC4", color= "blue") +
  labs(x = "Number of Workers", y = "Frequency", title = "Histogram of Number of Workers by Nesting Typ
  theme_minimal()

plot3
```

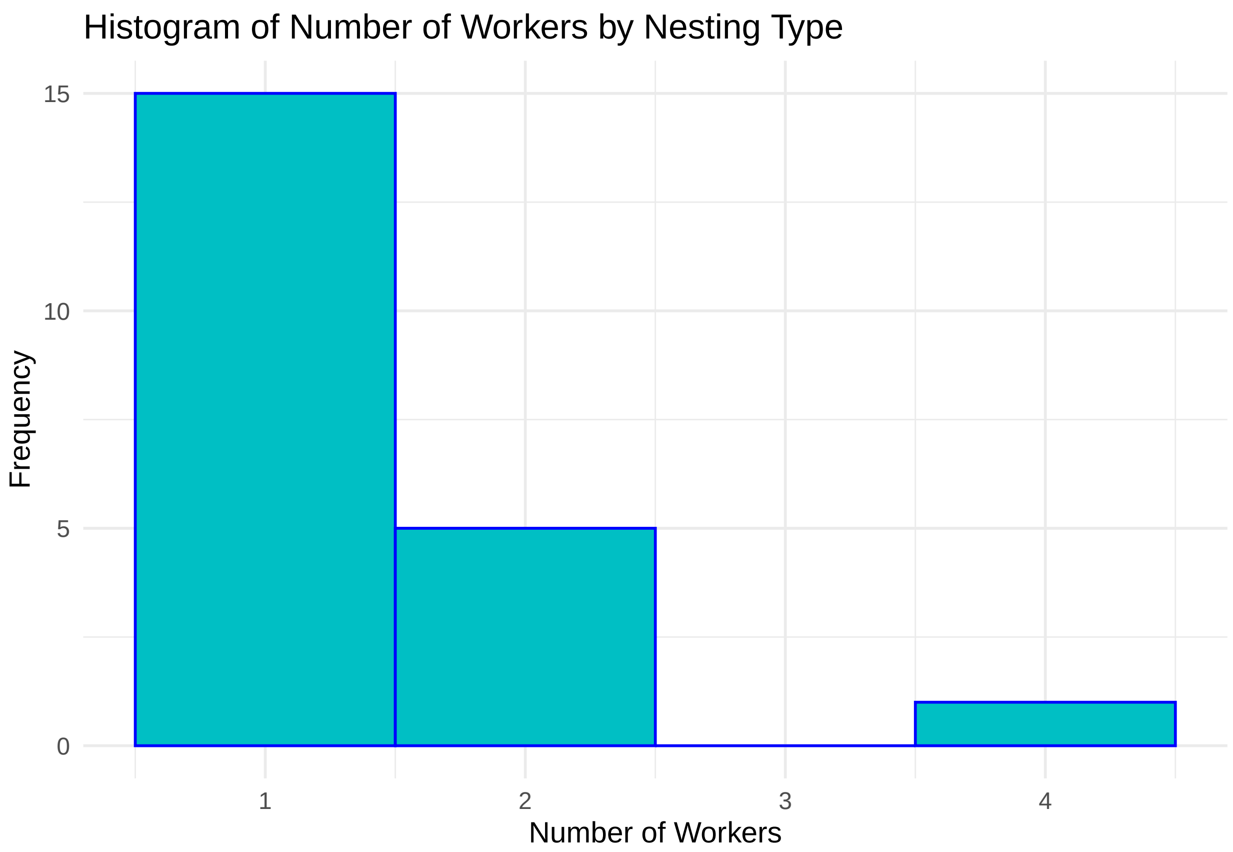


#### Session information - R version, packages, etc

Many thanks to all package creators!

```
## R version 4.3.3 (2024-02-29 ucrt)
## Platform: x86_64-w64-mingw32/x64 (64-bit)
## Running under: Windows 10 x64 (build 19045)
##
## Matrix products: default
##
##
## locale:
## [1] LC_COLLATE=English_United Kingdom.utf8
## [2] LC_CTYPE=English_United Kingdom.utf8
## [3] LC_MONETARY=English_United Kingdom.utf8
## [4] LC_NUMERIC=C
## [5] LC_TIME=English_United Kingdom.utf8
##
## time zone: Europe/Berlin
## tzcode source: internal
##
## attached base packages:
## [1] stats     graphics  grDevices utils     datasets  methods   base
##
```

```
## other attached packages:
##  [1] eoffice_0.2.2    ggfortify_0.4.17 coxme_2.2-20     bdsmatrix_1.3-7 
##  [5] survival_3.5-8   emmeans_1.10.1   DHARMa_0.4.6     car_3.1-2       
##  [9] carData_3.0-5    glmmTMB_1.1.9    ggplot2_3.5.1    readxl_1.4.3    
## 
## loaded via a namespace (and not attached):
##   [1] gridExtra_2.3           sandwich_3.1-0          rlang_1.1.3            
##   [4] magrittr_2.0.3          multcomp_1.4-25         compiler_4.3.3         
##   [7] mgcv_1.9-1              systemfonts_1.0.6       vctrs_0.6.5            
##  [10] devEMF_4.4-2            stringr_1.5.1           httpcode_0.3.0         
##  [13] pkgconfig_2.0.3         crayon_1.5.2            fastmap_1.1.1          
##  [16] magick_2.8.3            backports_1.4.1         labeling_0.4.3         
##  [19] utf8_1.2.4              promises_1.3.0          rmarkdown_2.26         
##  [22] nloptr_2.0.3            ragg_1.3.0              purrr_1.0.2            
##  [25] xfun_0.43               cachem_1.0.8            jsonlite_1.8.8         
##  [28] highr_0.10              later_1.3.2             uuid_1.2-0             
##  [31] broom_1.0.5             R6_2.5.1                R.devices_2.17.2       
##  [34] stringi_1.8.3           boot_1.3-29             cellranger_1.1.0       
##  [37] numDeriv_2016.8-1.1     estimability_1.5        Rcpp_1.0.12            
##  [40] knitr_1.46              zoo_1.8-12              base64enc_0.1-3        
##  [43] R.utils_2.12.3          httpuv_1.6.15           Matrix_1.6-5           
##  [46] splines_4.3.3           tidyselect_1.2.1        rstudioapi_0.16.0      
##  [49] abind_1.4-5             yaml_2.3.8              TMB_1.9.11             
##  [52] codetools_0.2-19        curl_5.2.1              lattice_0.22-5         
##  [55] tibble_3.2.1            shiny_1.8.1.1           withr_3.0.0            
##  [58] flextable_0.9.5         askpass_1.2.0           evaluate_0.23          
##  [61] gridGraphics_0.5-1      zip_2.3.1               xml2_1.3.6             
##  [64] pillar_1.9.0            plotly_4.10.4           generics_0.1.3         
##  [67] munsell_0.5.1           scales_1.3.0            minqa_1.2.6            
##  [70] xtable_1.8-4            glue_1.7.0              gdtools_0.3.7          
##  [73] lazyeval_0.2.2          tools_4.3.3             gfonts_0.2.0           
##  [76] data.table_1.15.4       lme4_1.1-35.3           fs_1.6.3               
##  [79] mvtnorm_1.2-4           grid_4.3.3              tidyr_1.3.1            
##  [82] colorspace_2.1-0        nlme_3.1-164            cli_3.6.2              
##  [85] rvg_0.3.3               textshaping_0.3.7       officer_0.6.5          
##  [88] fontBitstreamVera_0.1.1 fansi_1.0.6             viridisLite_0.4.2      
##  [91] dplyr_1.1.4             gtable_0.3.5            yulab.utils_0.1.4      
##  [94] R.methodsS3_1.8.2       digest_0.6.35           fontquiver_0.2.1       
##  [97] ggplotify_0.1.2         crul_1.4.2              TH.data_1.1-2          
## [100] farver_2.1.1            htmlwidgets_1.6.4       memoise_2.0.1          
## [103] htmltools_0.5.8.1       R.oo_1.26.0             lifecycle_1.0.4        
## [106] httr_1.4.7              mime_0.12               fontLiberation_0.1.0   
## [109] openssl_2.1.2           MASS_7.3-60.0.1
```
